# SER: an R package to compute environmental regime over a certain time period

**DOI:** 10.1101/2022.03.19.485011

**Authors:** Naicheng Wu, Kun Guo, Yi Zou, Fengzhi He, Tenna Riis

## Abstract

1. Environmental regime (or environmental legacy or historical legacy) is the environmental dynamic characteristics over a given (either long or short) time period, such as frequency of mean or extreme events and rate of change, which might be masked by using only contemporary variables.
2. We present SER, an R package for estimating environmental regimes for different environmental variables. Using the data included in the package, several examples are shown.
3. SER is suitable for any types of environmental variables e.g., nutrient concentration, light, dissolved oxygen. In addition, by changing the argument “days_bf”, it is possible to compute environmental regimes in any interested time period, such as days, months or years.
4. Our case study showed that inclusion of environmental regimes dramatically increased the explained variation of temporal β-diversity and its components. Environmental regimes, particularly in a given time period, are expected to advance the “environment - community” relationships in ecological studies. In addition, they can be implemented in other subjects, e.g., social science, socioeconomics, epidemiology, with important applied implications.

## 1. Introduction

A sound understanding of environment - community relations is a central topic in ecology. Scientists have been endeavoring to find suitable environmental variables or indices that have potential impacts on community compositions and distributions. Traditionally, snapshot contemporary environmental variables that were collected simultaneously with biological samples, such as *in situ* parameters, nutrient concentrations, are often employed. However, this neglects the fact that biological community responds not only to contemporary environmental condition but also to historic environmental (also called historic legacy) characteristics (Fig. 1) (Su *et al*. 2022). For example, Oliveira, Lima-Junior and Bini (2020) found that current environmental variables were weak predictors of fish community structure, and that the predictive power substantially increased when using dataset obtained in a previous time period. In response, new indices that integrate long-term environmental records were proposed. For instance, hydrologic indicators for characterizing streamflow regimes (i.e., flow regimes) using long-term flow records have been developed to represent biologically relevant streamflow attributes (Olden & Poff 2003). Another example is the nineteen standard bioclimatic indices, which integrate climate data from 1970-2000 (available in WorldClim 2 database; Fick & Hijmans 2017). In addition, historical legacies (i.e., past climate and geography: temperature anomaly during the quaternary period, past temperature trend, past precipitation trend, past climate-change velocity, basin median latitude and the endorheic/exorheic status of the river) were computed and used to explore their roles to the functional diversity of global freshwater fishes (Su *et al*. 2022). The results showed that the historical legacies significantly imprinted the functional dispersion and functional identity patterns.

**Fig. 1.**
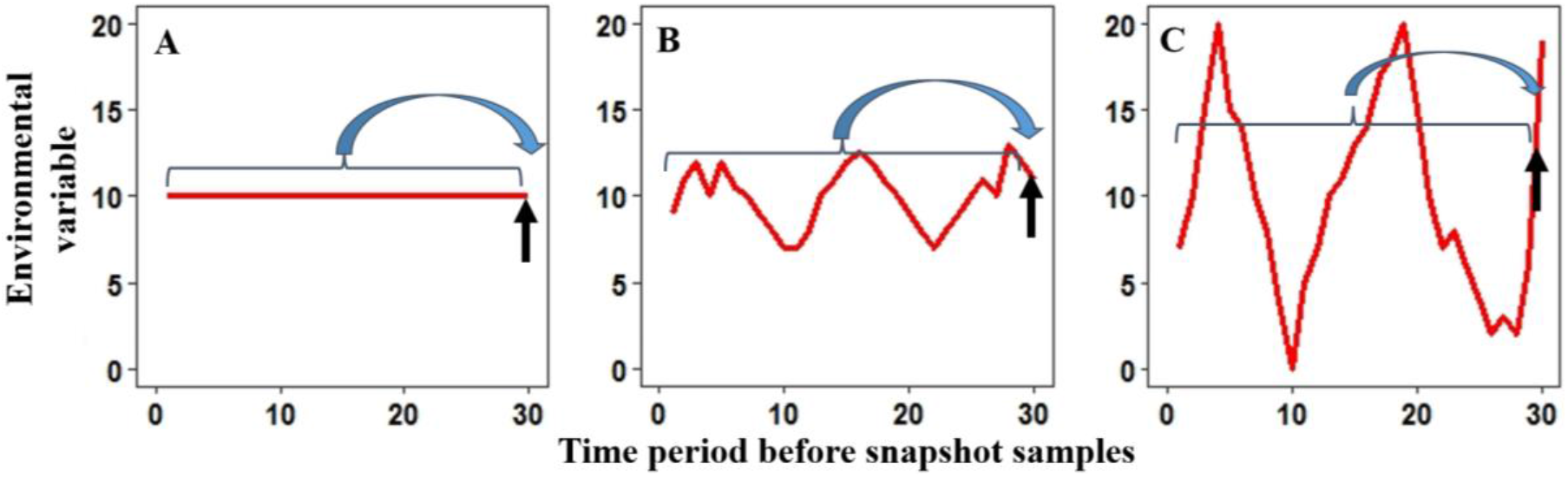
Schematic figure showing 3 scenarios of enviromental conditions over a certain time period (within curly brackets) before snapshot samples (indicated by black arrows). Scenario A: environmental variable being constant; B: weak fluctuations in the environmental variable; and C: strong fluctuations in the environmental variable over the whole time period (i.e. from time 0 to 30). Commonly used indices such as instant value, simple mean or median values do not sufficiently represent the enviroment regimes/fluctuations prior to the sampling date in scenario B and C.

The utility of these historical environmental regime indices has resulted in tremendous applications in ecological research and also other lines of research (e.g., Tonkin *et al*. 2018; Nguyen *et al*. 2021; Tornés *et al*. 2021; De Pauw *et al*. 2022; Su *et al*. 2022; Xu *et al*. 2022). However, there are several constraints to the currently used historical environmental regime indices: 1) the current available indices are limited to hydroclimatic variables, such as flow, temperature, precipitation. There is no available R package to calculate these indices of other data such as pH, turbidity, dissolved oxygen, chlorophyll a, and other biotic data (e.g., animal feeding or movement); 2) these indices are mostly based on long-term intervals, e.g., 30 years for bioclimatic variables. Given that some organisms, particularly microorganisms, may show quick responses to environmental changes, the aforementioned indices might fail to link with biotic changes, and shorter time period may be more relevant. In addition, different organisms (e.g., algae, macroinvertebrates, fish, macrophytes, or even terrestrial plants) have distinct extent of response to historical environmental regimes. For instance, recent studies found that flow regimes over a short-term period (e.g., 7 or 14 days) played a vital role in riverine algae and biofilm communities (Wu *et al*. 2018; Qu *et al*. 2019; Guo *et al*. 2020; Guo *et al*. 2021). In contrast, macroinvertebrate and fish communities may show a good response to environmental changes over a longer time period (e.g., 4 weeks, 1 year) (Schneider & Petrin 2017). Therefore, to differentiate the distinct responses of different organisms, we should derive community specific indices that describe environmental patterns over relevant time periods. Unfortunately, no R package so far provides a function to calculate indices over a required time period.

Prompted by the importance of environmental characteristics over a certain time period and their research scarcity in this field, we here propose a new term of “environmental regime” (or environmental pattern or environmental legacy or historical legacy). Unlike the traditional environmental variables, these new environmental regime indices are defined as the environmental dynamic characteristics during a given (either long or short) time period, which might be masked by using contemporary environmental variables or simple average or median values (Fig. 1). With the facilitation of science and technology, more and more high frequently (daily, sub-daily, hourly, or even finer scale) measured environmental variables (e.g., nutrient concentration, dissolved oxygen) are available nowadays. An increasing number of studies have been using data from high-frequency measurements, e.g. limnology (Meinson *et al*. 2015) or soil greenhouse gas fluxes (Courtois *et al*. 2019). These data provide scientists a chance to explore research questions at time scales that were not possible earlier. Further, high-frequency data allow computing environmental regimes that can be potential variables to increase explained variation of biological communities (e.g., Wu *et al*. 2019; Guo *et al*. 2020; Wijewardene *et al*. 2021a). Nevertheless, an R package or function that can be easily used to calculate short-term environmental regime is still missing.

## 2. The SER package: Short-period Environmental Regime

In total, 11 elementary indices that focus on variations of environmental factors in a given short-period were developed (Table 1). These indices, inspired by Olden & Poff, (2003), elucidate three aspects i.e., the magnitude, the frequency, and the rate of change of environmental variables in a given time period. The magnitude contains four categories: mean, median, coefficient of variation and skewness of the variables in a given time period before the snapshot sampling; frequency demonstrates the number of environmental low or high pulses in a given time period before the snapshot sampling; rate of change how fast the environmental variable changed (i.e., positive or negative change) within the given time period before the snapshot sampling.

**Table 1.** Definition of the 11 elementary indices for environmental regimes.

| Code | Unit | Definition |
| --- | --- | --- |
| **Magnitude of environmental data** |  |  |
| MA1 | Depends | Mean of the environmental variable in a given time period (N) |
| MA2 | Depends | Median of the environmental variable in a given time period (N) |
| MA3 | % | Coefficient of variation in a given time period (N) |
| MA4 | - | Skewness of the environmental variable for a given time period (N) |
| **Frequency of environmental data** |  |  |
| ML1 | Temporal unit | Number of low pulse events in a given time period (N):<br>numbers of occurrences where the magnitude of an environmental variable drops below the 25th percentile of all values for the given time period. |
| MH1 | Temporal unit | Number of high pulse events in a given time period (N):<br>numbers of occurrences where the magnitude of an environmental variable is above the 75th percentile of all values for the given time period. |
| EL1 | Temporal unit | Number of extreme low pulse events in a given time period (N):<br>numbers of occurrences where the magnitude of an environmental variable drops below the 10th percentile of all values for the time period. |
| EH1 | Temporal unit | Number of extreme high pulse events in a given time period (N):<br>numbers of occurrences where the magnitude of an environmental variable is above the 90th percentile of all values for the time period. |
| **Rate of change of environmental data** |  |  |
| RC | Depends | Rate of change in a given time period (N). |
| RH1 | Temporal unit | Numbers of events with rising change in a given time period (N), i.e., numbers of positive change |
| RL1 | Temporal unit | Numbers of events with declining change in a given time period (N), i.e., numbers of negative change |

Depends: the unit depends on the variables; - : no unit; Temporal unit: the given temporal units, e.g., days, months or years. N can be defined as any interested time period, by default N=BetwSamT (days between two successive sampling dates). Currently, the package is developed with regarding days, months and years as the interested period, more features are under development.

### Package overview

The SER package contains one main function *SER* and two data files i.e., *hydro_df* and *sample_date*. The two data files are derived from (Guo *et al*., 2020) and are used to illustrate how the main function works. The *hydro_df* is a data frame that contains daily discharge in a stream, while *sample_date* is a vector containing 13 dates, first of which is the date when experiment was initialized while the rest 12 are snapshot biological sampling dates.

The SER package is written in R (R Core Team 2021) and require a standard installation of R and the ‘tidyverse’ and ‘lubridate’ packages. The development version of SER is hosted on GitHub at https://github.com/kun-ecology/SER. The package can be installed via below code:

~~~
if(!require(devtools)) install.packages(“devtools”)
devtools::install_github(“https://github.com/kun-ecology/SER“, build_vignettes= TRUE)
~~~

### Example analyses

As an example, the embedded data are used to illustrate how SER works with discharge data. By default, days between two successive sampling dates was used as the focal short-period. The R codes are shown as below:

~~~
# load required packages
library(tidyverse)
library(lubridate)
library(SER)
# inspect the discharge data
str(hydro_df)
# make sure that data column (in this case Discharge) is “Env”
names(hydro_df)[2] <- “Env”
# inspect the sample dates
str(sample_date)
# calculate short-period hydrological indices
hydro_ser <- SER(hydro_df, sample_date, days_bf = NULL)
str(hydro_ser)
~~~

The function SER returned a data frame, in which, the first column ‘SampleDate’ contains the 12 biological sampling dates, while the rest 11 columns represent the short-period environmental regimes, i.e., short-period hydrological indices for each sampling date (Fig. 2). The indices’ names were constructed as the combination of short time period and names of the elementary indices, for example, BetwSamT.MA1 and BetwSamT.RC stand for mean of the daily average flow and mean rate of change in days between two successive sampling days respectively.

**Fig. 2.**
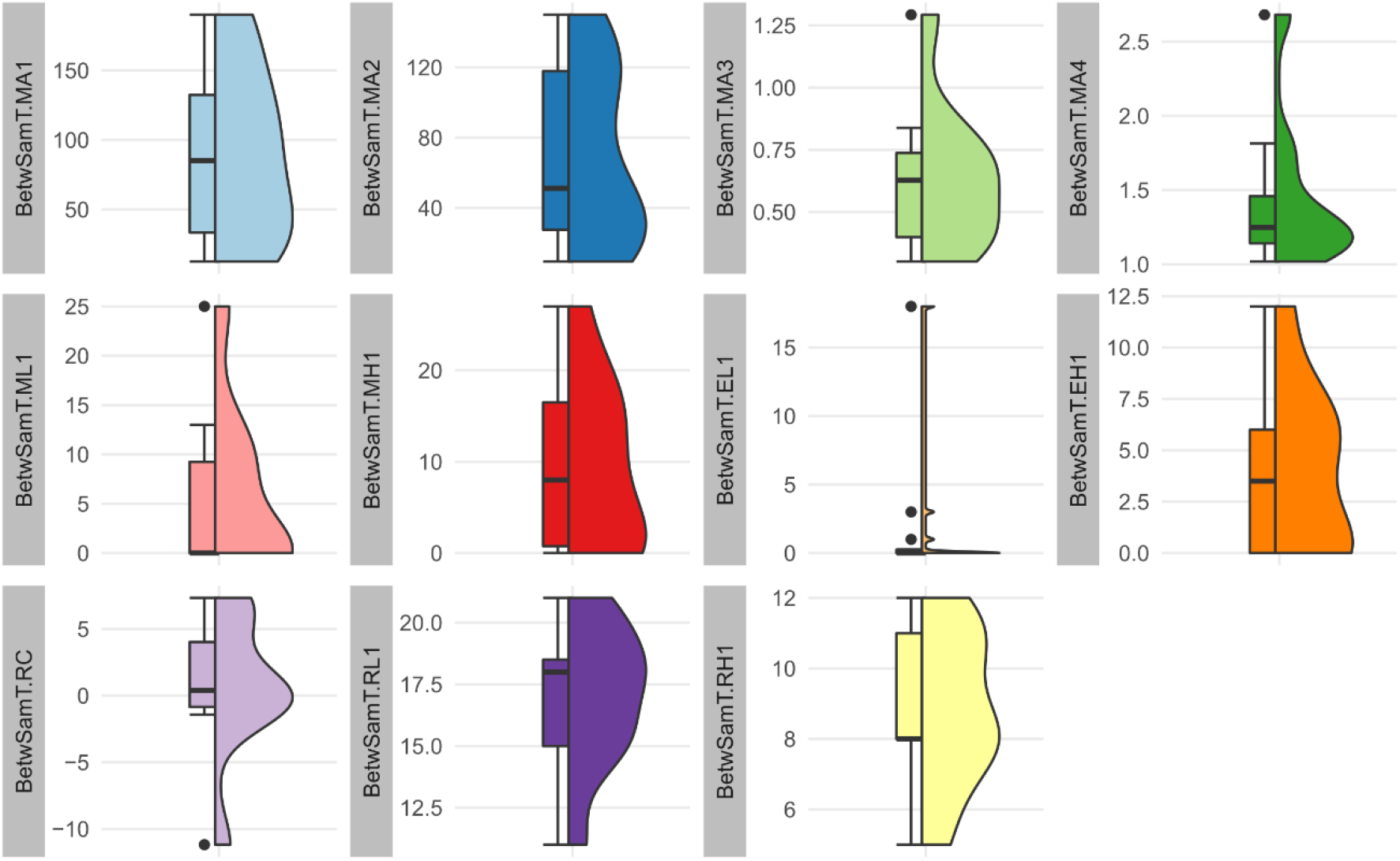
Boxplots (median, first, and third quantiles) and violin plots illustrate the sample density for the 11 short-period hydrological indices described in Table 1.

## 3. A case study: Environmental regimes play an important role for taxonomic and functional temporal β-diversity of riverine diatoms

To examine whether inclusion of environmental regimes advance our understanding of environment-biota relationships, daily samples of riverine diatom communities over a 1-year period were collected at a German lowland catchment (Wu *et al*. 2022a). Concurrently, three categories of abiotic factors were obtained: a) hydrological variables (Hyd) included daily discharge (Q), discharge skewness (Sk), precipitation (Prec), and antecedent precipitation index (API); b) metal ions (Met) contained six parameters (Cl^-^, K^+^, Ca^+^, Na^+^, Mg^2+^ and Si^2+^); and c) nutrients (Nut) included ammonium-nitrogen (NH_4_-N), nitrate-nitrogen (NO_3_-N), orthophosphate (PO_4_-P), and sulphate (SO_4_^2-^). In addition, environmental regimes of both flow (using Q) and nutrient (using NH_4_-N, NO_3_-N, PO_4_-P and SO_4_^2-^) were computed with *SER* package. Therefore, we have two extra abiotic factors: Hyd+ (i.e., hydrology + flow regimes), and Nut+ (i.e., nutrient + nutrient regimes). Furthermore, both taxonomic and functional temporal β-diversity of riverine diatoms were computed and then decomposed into turnover and nestedness components (for details see Wu *et al*. 2022a).

Using distance-based redundancy analysis (db-RDA) and variation partitioning analysis (VPA), we investigated the relationships between abiotic factors and temporal β-diversity of riverine diatoms (for details see Wu et al. 2022). To detect the role of environmental regimes in explaining the variation of temporal β-diversity and its components, we compared the explained variations between without and with environmental regimes. VPA results demonstrated that addition of environmental regimes (i.e., flow and nutrient regimes) clearly increased the explained variations of both taxonomic and functional temporal β-diversity and its components (Fig. 3 and 4). Specifically, taxonomic total β-diversity, turnover and nestedness increased by 3.0%, 4.9% and 15.5%, respectively, while functional total β-diversity and its components increased by 13.3%, 4.6% and 12.2%, respectively. Interestingly, inclusion of flow regimes (i.e., Hyd+) played a less important role in taxonomic temporal β-diversity than functional temporal β-diversity. In contrast, addition of nutrient regimes (i.e., Nut+) explained more variations in taxonomic temporal β-diversity than functional temporal β-diversity (Fig. 3). Regardless of the potential reasons, which warrant further investigations, these results supported our hypothesis that addition of environmental regimes could dramatically advance our understanding of environment-biota relationships.

**Fig. 3.**
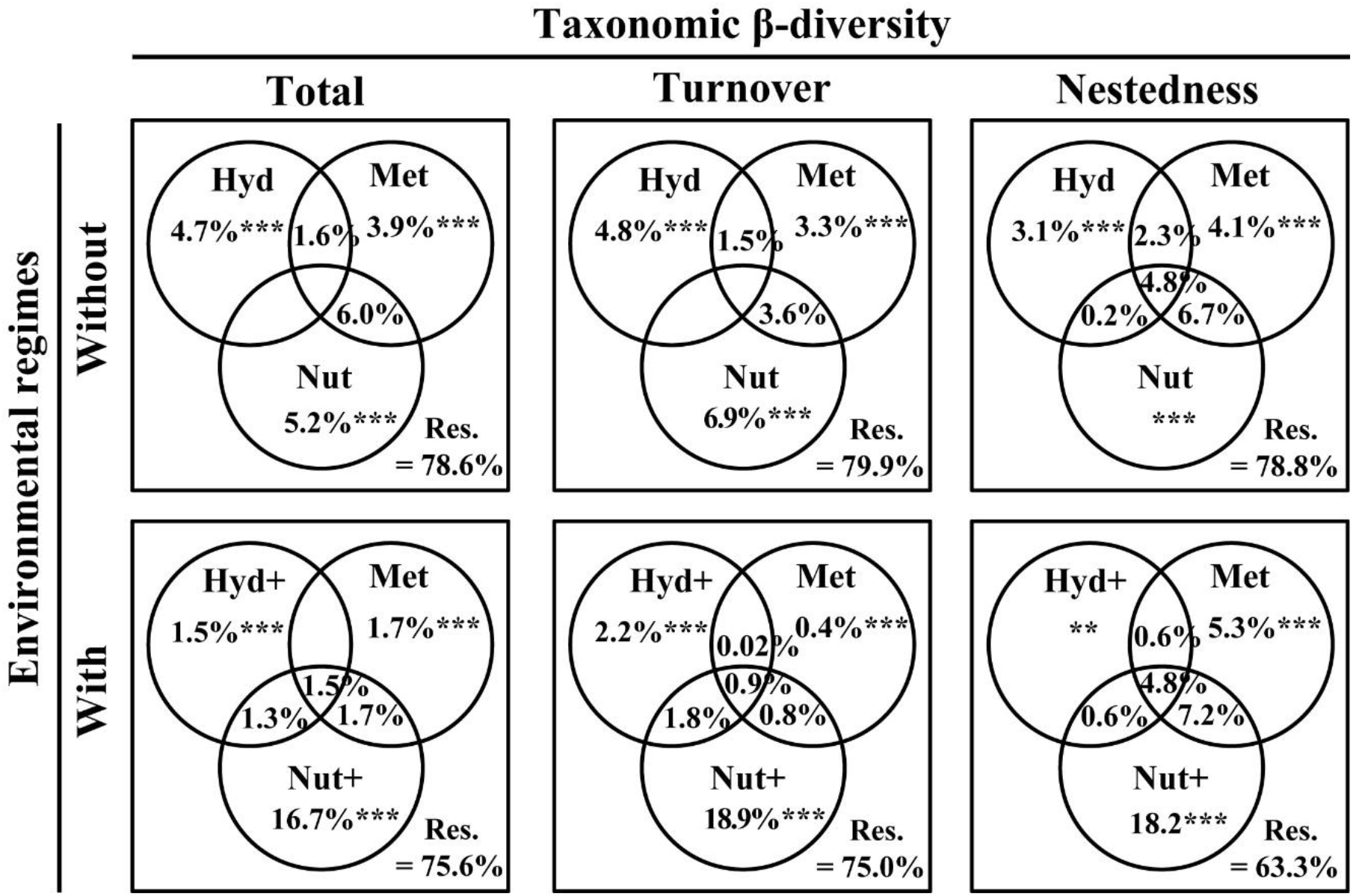
Comparison (between without and with environmental regimes) of the explained variations to taxonomic temporal β-diversity components of riverine diatoms. Hyd = hydrology without flow regimes; Met= metal ions; Nut= nutrients; Hyd+ = hydrology with flow regimes; Nut+= nutrients with nutrient regimes. The adjusted R^2^ is shown. *** p < 0.001, ** p < 0.01, * p < 0.05. The figure was modified from Wu *et al*. (2022).

**Fig. 4.**
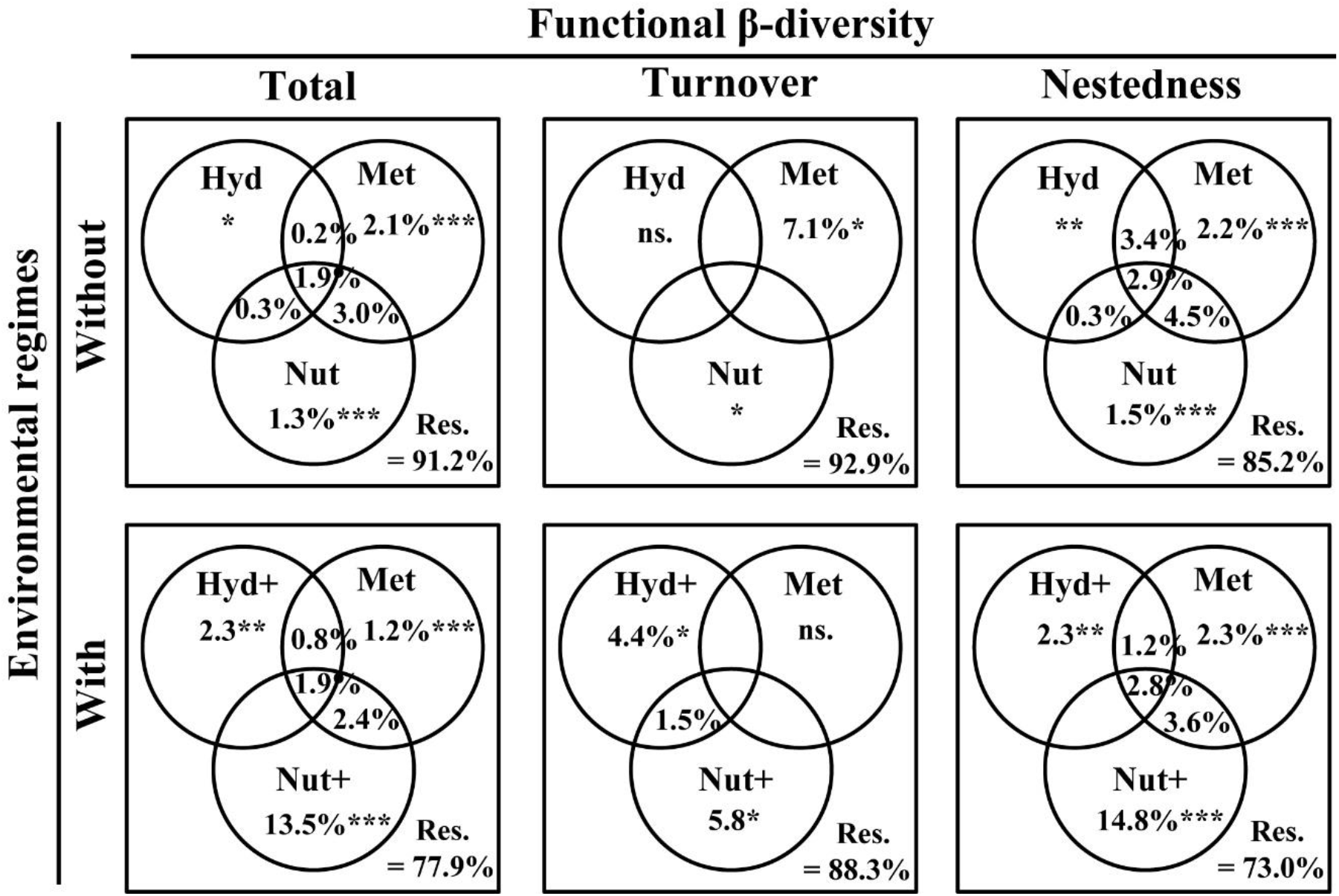
Comparison (between without and with environmental regimes) of the explained variations to functional temporal β-diversity components of riverine diatoms. Hyd = hydrology without flow regimes; Met= metal ions; Nut= nutrients; Hyd+ = hydrology with flow regimes; Nut+= nutrients with nutrient regimes. The adjusted R^2^ is shown. *** p < 0.001, ** p < 0.01, * p < 0.05. The figure was modified from Wu *et al*. (2022).

## 4. Conclusion and remarks

SER is an important tool to facilitate calculation of environmental regimes over a given time period. As a holistic term, it is suitable for any types of environmental or biotic parameters, such as nutrient concentration, pH, conductivity, light, dissolved oxygen, chlorophyll a. Furthermore, by changing the argument “days_bf”, it is possible to compute environmental regime in any given time period, such as months or years, as long as the records are measured in a corresponding manner.

Being a completely open source tool, it’s open for further extension and examination. We envisage that SER is greatly helpful in both basic and applied ecological studies from mesocosm experiments to field surveys. On one side, environmental regimes (e.g., thermal, nutrient, flow), particularly short-term environmental regimes, can be robust variables in understanding the “community-environment” relationships of different organisms in various ecosystems (e.g., aquatic, forest, terrestrial ecosystems), being complementary predictors for model simulation and prediction. A recent study found that severe changes in the thermal regimes of Austrian rivers under climate change reinforced physiological stress and supported the emergence of diseases for brown trout (Borgwardt *et al*. 2020). On the other side, exploring responses of different organisms to environmental regimes shifts can be used for management and policy making. For instance, by exploring the relationships between the occurrence of cyanobacterial blooms and water-level regimes, management of water-level can be a potential mitigation strategy for cyanobacterial blooms (Bakker & Hilt 2016). In addition, we would like to emphasize SER’s potential in experimental biology or mesocosm experiments, which often last for a relative short period but could have high-frequency measured data, e.g., temperature, light. High-frequency data (at 15-minute interval) of light and water temperature were measured in a microcosm study, and the results indicated light and temperature emerged as significant variables on phytoplankton community attributes (Wijewardene *et al*. 2021b).

To a broad extent, environmental regimes can be used in other subjects, e.g., social sciences, socioeconomics, epidemiology. As an example, a recent study (Wu *et al*. 2022b) found that increasing temperature variability (calculated as the standard deviation of the average of the same and previous days’ minimum and maximum temperatures) has caused a higher human heat-related mortality. Another example showed that shift in a temperature regime caused by climate changes may facilitate a pathogen’s survival, development and spillover, and have an effect on transmission chains. Pandemic forecasting models (such as COVID-19) were recommended to integrate these effects, alongside human behavior and awareness (Rodó et al. 2021). A third example is about crop yield in relation to weather regimes. Altering temperature and rainfall regimes, such as unusually cool and wet spring, is reducing global production of staples (e.g., rice, wheat), while, in contrast, some more drought-tolerant crops (e.g., sorghum) have benefited (Ray et al. 2019). Developing an empirical model linking crop yield to weather regimes may inform local people with proper crops under future climate scenarios.

## Acknowledgements

This study was supported financially by the Humboldt fellowship for experienced researcher and the Starting Grants of Ningbo University (Nos. 421999292, 422110123).

## Author’s contribution

K.G. developed the package; N.W. and K.G. wrote the manuscript; K.G. tested all the codes and the examples; Y.Z., F.H. and T.R. revised and improved the manuscript. All authors have approved the final version to be published.

## Conflict of interest

The authors declare no conflict of interest.

## Data availability statement

The SER package can be downloaded from GitHub (https://github.com/kun-ecology/SER). An online tutorial is available for this package on the same GitHub repository. SER depends on two existing R packages: tidyverse and lubridate.

## Notes

### Competing Interest Statement

The authors have declared no competing interest.

